# Characterization of Soybean Vein Necrosis Virus (SVNV) Proteins: Sequence Analysis of Field Strains and Comparison of Localization Patterns in Differing Cell Types

**DOI:** 10.1101/2024.11.26.625476

**Authors:** Abdelaal H.A. Shehata, Michael A. Mayfield, Edward J. Sikora, Kathleen M. Martin

**Affiliations:** Department of Entomology and Plant Pathology, Auburn University, Auburn AL

**Keywords:** Soybean vein necrosis virus, orthotospovirus, co-localization, cell death, strain sequencing, and *Neohydatothrips variabilis*

## Abstract

*Soybean vein necrosis virus* (SVNV) is a persistent, propagative, ambisense-ssRNA virus in the genus Orthotospovirus transmitted by *Nehydratothrips variabilis*. In this study, five open reading frames (ORFs) (N, NSs, NSm, G_N_, and G_C_) were fused in frame to GFP for use in both plant and insect cells. N and NSs localize in plants to the cell periphery and nucleus. NSm causes cell death to plant cells but not in insect cells where cytoplasmic localization is seen. G_N_ and G_C_ glycoproteins localize to the membranes and accumulate with more cytoplasmic localization seen in insect cells. To understand SVNV in the field, we collected 33 samples expressing symptoms of SVNV. The N, NSs, and NSm ORFs were sequenced and amino acid mutations in each gene were seen. The findings of this study capture sequence changes which have occurred over the past fifteen years compared to the localization of proteins never before conducted.

## Introduction

Soybean (*Glycine max*) is a globally significant crop renowned for its nutritional value and versatility. Originating in East Asia, soybeans are now cultivated worldwide, particularly in regions with temperate climates (Hymowitz, 1970). This leguminous plant plays a pivotal role in agriculture serving as a primary source of protein and oil for both human consumption and livestock feed (Wilcox, and Shibles 2001). Soybeans are also prized for their adaptability to diverse agricultural practices and their ability to fix nitrogen in the soil through symbiotic relationships with beneficial bacteria (Herridge et al., 2008). With a multitude of applications ranging from food products to industrial uses, soybean remains a cornerstone of modern agriculture and food security initiatives globally (Boehlje and Brorsen, 2009). Due to all these features, soybean is a commonly planted crop among farmers in Alabama and is ranked as the fourth most grown crop in the state with a production of 14.55 million bushels (870 million lbs) in 2022 with an estimated value of 206.6 million dollars (USDA 2022).

Soybean vein necrosis virus (SVNV) was first discovered in Tennessee in 2008 (Tzanetakis et al., 2009). Symptoms of the virus include yellowing and brown necrotic tissues along the major veins of the upper and lower leaf surface which results in a scorched appearance of damaged leaves (Conner et al., 2013). SVNV also impacts the growth, seed quality, and oil content of soybean (Anderson et al., 2017 and Groves et al., 2016) and is transmitted in a persistent, circulative manner by various species of thrips. Soybean thrips, *Neohydatothrips variablilis*, are the most efficient vector of SVNV, with transmission rates ranging from 67-100% (Zhou and Tzanetakis, 2013). However, other species of thrips such as *Frankliniella fusca*, *Frankliniella tritici*, and *Frankliniella schultzei* can also transmit at very low rates (Han et al., 2019). As with all Orthotospoviruses, SVNV can only be acquired by thrips during its first or second instar larval stages for viral transmission to occur as adults (Han et al., 2019) In Alabama, SVNV was first reported in 2012 (Conner et al., 2013) and between 2013 to 2016 an increase in the average incidence of SVNV on soybean from 31.8% to 82.6% was reported (Chitturi et al., 2018). Since the discovery of SVNV, it has been confirmed in 22 states (Tzanetakis et al., 2009, Conner et al., 2013).

SVNV is an ambisense, ssRNA virus belonging to the genus *Orthotospovirus* in the family *Bunyviridae*. The SVNV genome consists of three genomic segments encoding five ORFs (Zhou et al., 2011). The S segment has two ORFs, the non-structural small component (NSs, 1323 bp, silencing suppressor) and the nucleocapsid (N, 834 bp, encases the viral RNA) with a total length of 2,603 bp. The M segment contains two ORFs: the non-structural medium component (NSm, 951 bp, movement protein) and the combined sequence for the glycoproteins (G_N_ and G_C_) with a total length of 4,955 bp. The last genomic segment (L) is the largest; it only contains a single ORF encoding an RNA-dependent RNA-polymerase (RdRp) with a length of 9,010 bp (Zhou et al., 2011). Each of the structural proteins is responsible for the formation of the completed virion, with N encasing the viral RNA, interacting with the glycoproteins embedded in host membranes to bud through these membranes to form the completed virion (Snippe et al., 2005 and Tsompana and Moyer 2008). G_N_ and G_C_ are located on the outer surface of the virion and are expected to be required for transmission by their insect vector, *Neohydatothrips variabilis* (Whitfield et al., 2004).

To understand SVNV better, the population structure and diversity of SVNV was studied among different states by Zhou and Tzanetakis (2013). Between 2008 and 2012, 37 SVNV isolates were collected from Arkansas, Delaware, Illinois, Kansas, Maryland, Mississippi, Missouri, and Tennessee (Zhou and Tzanetakis 2013). The RNA and protein alignments of the nucleocapsid protein of the 37 isolates were constructed with another 11 SVNV isolates that were collected from Kentucky and Tennessee (Khatabi et al., 2012). The RNA and protein comparisons showed identities of 98 to 100% and 97 to 100% subsequently, which suggest that SVNV has a relatively homogeneous population structure within the geographic states studied (Zhou and Tzanetakis 2013). Moreover, the phylogenetic analysis did not confirm a distinct geographical pattern in isolate clustering which also indicates that there is no significant diversity observed within or among populations in the states studied (Zhou and Tzanetakis 2013).

In this study, we are interested in understanding the relationship of SVNV with other orthotospoviruses and determining why SVNV is found in a distinct clade in the phylogenic studies that were conducted previously (de Oliveira et al., 2012, Oliver and Whitfield 2016, and Kormelink et al., 2021). The hypothesis guiding this study is that the localization pattern of SVNV proteins is likely to be similar, but not identical, to other characterized orthotospoviruses and that the sequence of SVNV has likely undergone changes since the original sequencing study was conducted over ten years ago. To address these questions, we localized and co-localized SVNV proteins in plant and insect cells and documented differences in SVNV protein localization compared to other characterized species. We also investigated genetic changes have occurred in SVNV from Alabama indicating possible adaptations over time compared to the sequencing conducted previously in other states. These findings will further our understanding of this important virus and assist in continuing to develop management options directed at this threat to soybean production in Alabama and across the United States.

## Materials and Methods

### Plant Growth and Maintenance

Wildtype, and transgenic marker *N. benthamiana* plants expressing red fluorescent protein fused either the endoplasmic reticulum (RFP-ER) and histone 2B (RFP-H2B) (Martin et al., 2009) were cultivated in a growth chamber maintained at 25°C with a light intensity of 300 µM for 14 hours of daylight, followed by a 10-hour dark period. At two weeks of age, the seedlings were transferred to individual containers. Subsequently, the plants were allowed to grow for one to two more weeks until their leaves reached the size of greater than 24 mm before use in infiltrations for microscopy.

### The source of SVNV strain and the amplification of the five ORFs of SVNV

To test the protein localization and co-localization of SVNV, the reference genome of the SVNV strain of Tennessee (accession: GCF_004789395.1) was used to clone five ORFs of SVNV (N, NSs, NSm, G_N_, and G_C_). One microliter of the gene template (50 ng) was combined with specific primers containing CACC before the start codon to amplify each gene (Table 1). L was not included due to its’ extreme size (9,010 bp). The amplifications were carried out using Phusion High-Fidelity Enzyme using the manufacturer’s protocol (Thermo Fisher Scientific, Waltham MA) with adjusting the primer annealing temperature to our specific primers (Table 1). The expected band size for each gene (Table 1) was purified and recombined into pENTR D-TOPO following the manufacturer’s protocol (Invitrogen^TM^). For the glycoproteins (G_N_ and G_C_), reverse primers (G_N_-S-R and G_C_-S-R) (Table 1) were also designed to trim the transmembrane domain (TM) (Figure 1) mapped using the methods described below. Successful cloning was confirmed using PCR with the M13-F and M13-R primers and sent for Sangar sequencing.

**Table 1.**
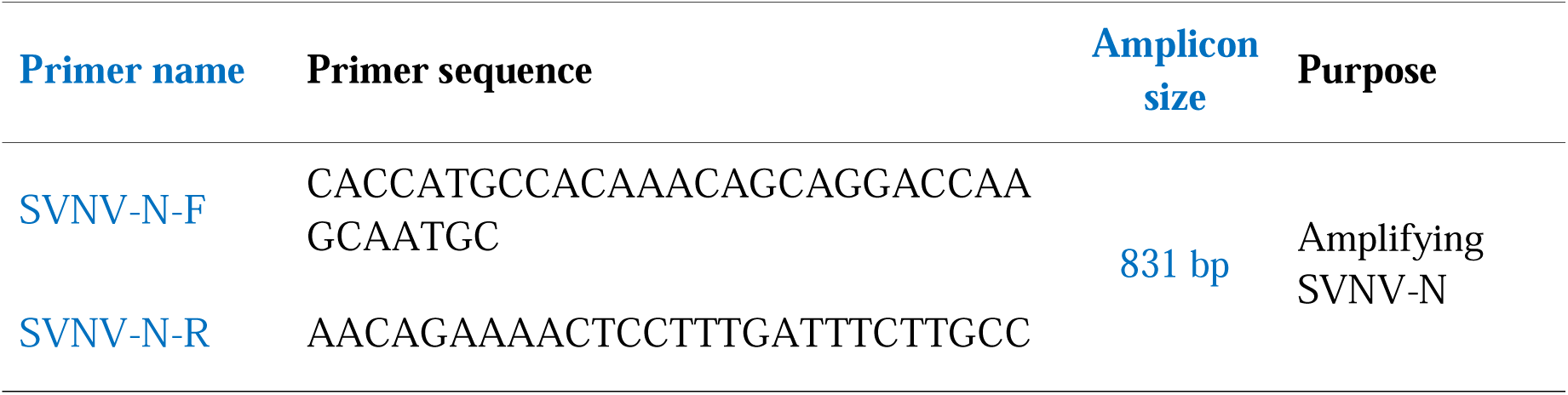
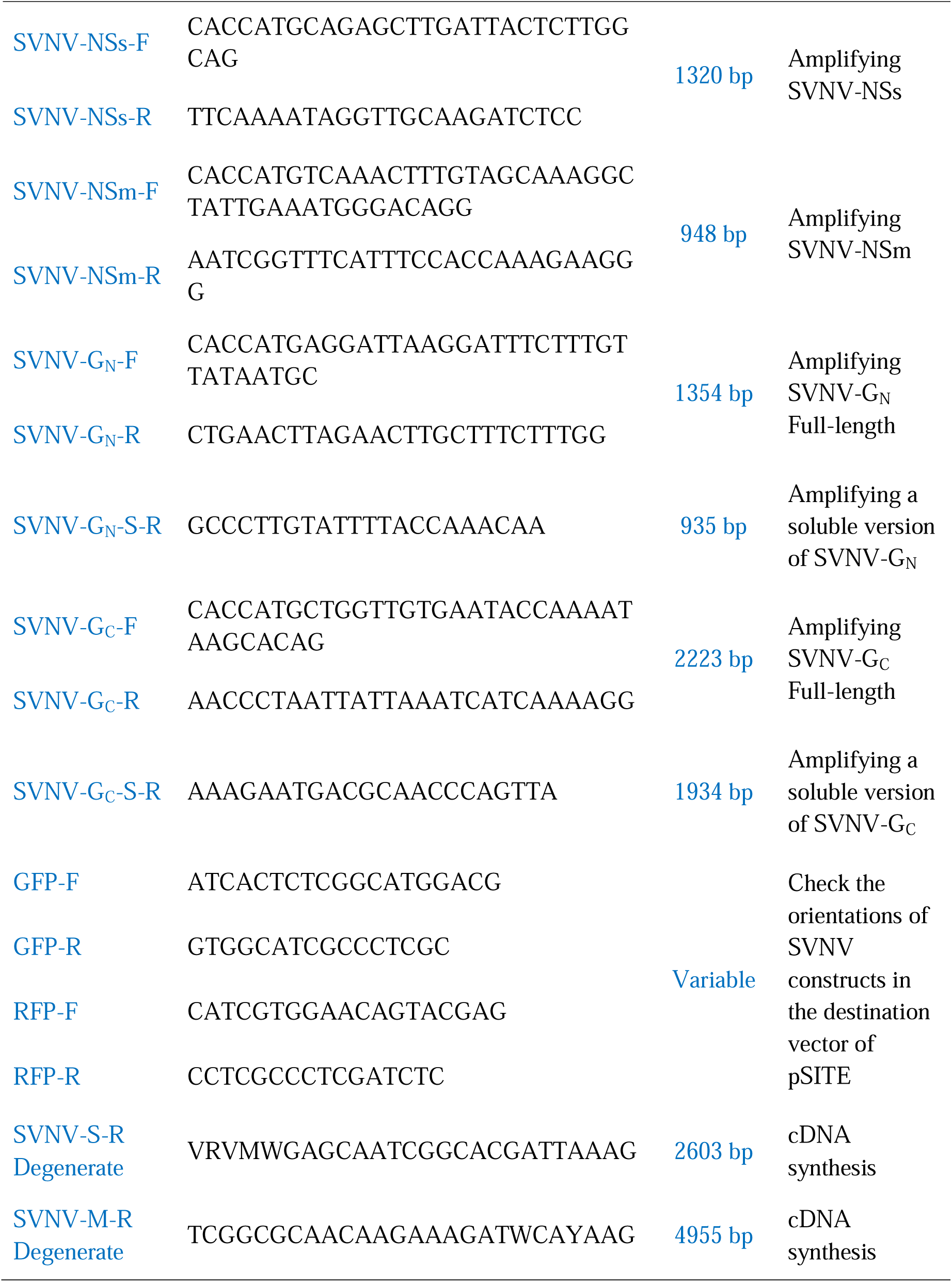
The specific primers that were used in amplifying SVNV genes, The specific reverse primers of SVNV-GN and GC to produce a soluble version of the glycoproteins (GN and GC), The GFP/RFP forward and reverse primers that were used with the appropriate SVNV gene-specific primer (Forward or reverse) to check the orientation of the constructs in pSITE-2NA, pSITE-2CA, pSITE-4NA, and pSITE-4CA.

**Figure 1:**
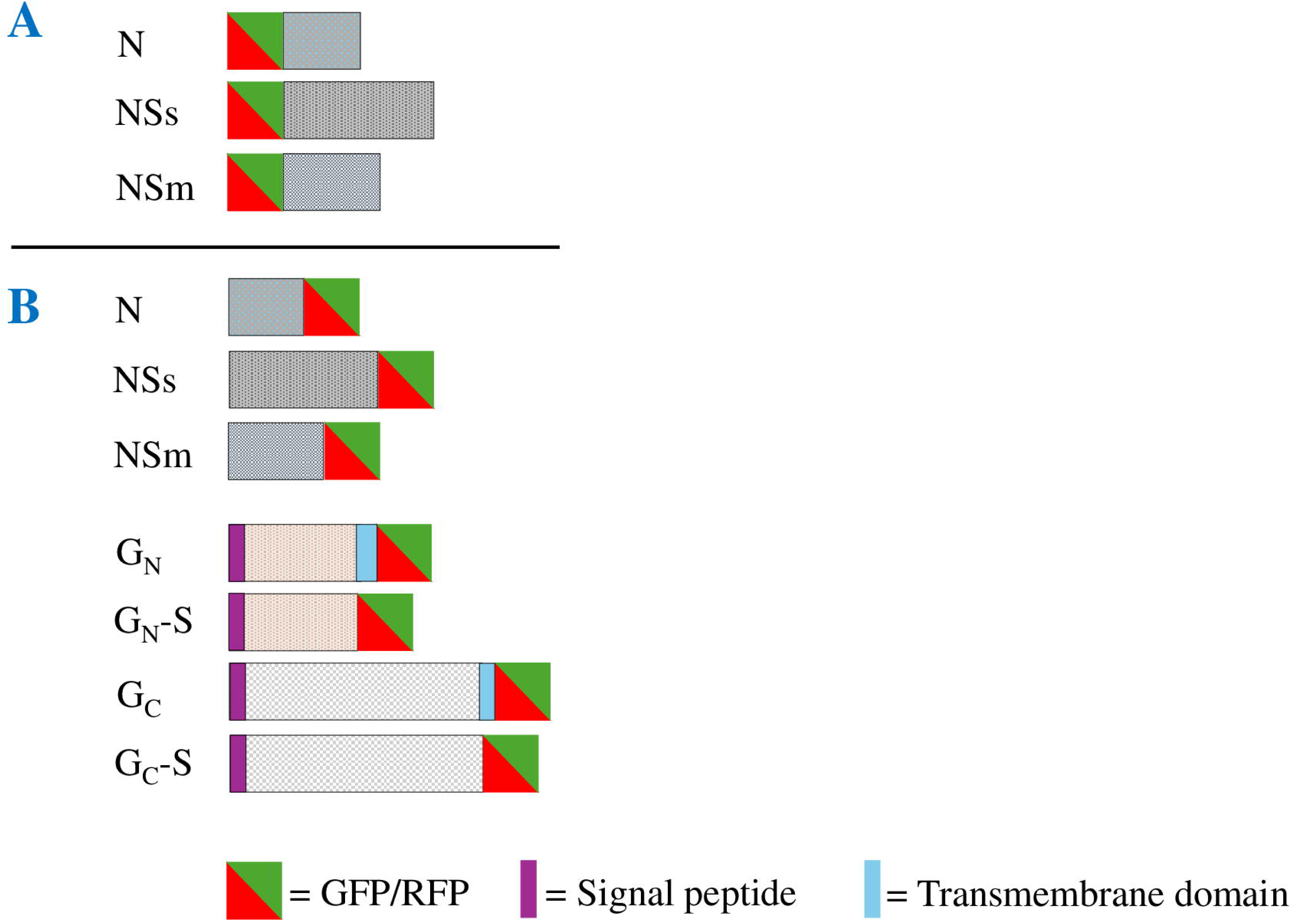
Graphic representation of the fused Soybean vein necrosis virus constructs in the pSITE destination vector used in fluorescence localization. A shows the constructs that were fused in the C terminal of GFP or RFP. B shows the constructs that were fused in the N terminal of GFP or RFP.

### Domain analysis of SVNV

Protein domains for SVNV were identified using My Hits (Swiss Institute of Bioinformatics) Motif Scan. Parameters were set to include “hamap - HAMAP profiles”, “pat - PROSITE patterns”, “freq_pat - frequent match producers for “PROSITE patterns”, “pre - More profiles”, and “ prf - PROSITE profiles. The signal peptide and the transmembrane domains of SVNV were determined and analyzed using TMHMM 2.0 and InterPro (https://services.healthtech.dtu.dk/services/TMHMM-2.0/ & https://www.ebi.ac.uk/interpro/). (Krogh et al., 2001, Sonnhammer et al.,1998, and Paysan-Lafosse et al., 2022) The Nuclear Localization Signals (NLSs) of SVNV proteins were identified using the NLS domain mapper (https://nls-mapper.iab.keio.ac.jp/cgi-bin/NLS_Mapper_form.cgi) (Kosugi et al., 2008, Kosugi et al., 2009a, and Kosugi et al., 2009b). These domains and their positions for SVNV were compiled into tables using Microsoft Excel (Build 17231.20236 Click-to-Run). *Visualization of SVNV ORFs in plant cells*

After the confirmation of SVNV ORFs in pENTR-d-Topo, LR reactions were carried out with pSITE-2CA, pSITE-2NA, pSITE-4CA, and pSITE-4NA (Chakrabarty et al., 2007; Martin et al., 2009) following the manufacturer’s recommendations (Invitrogen^TM^) (Figure 1). The positive colonies (in both orientations) were confirmed by PCR using primers for the fluor and the gene in question (Table 1). After the SVNV constructs were confirmed, they were transformed into *Agrobacterium tumefaciens* LBA 4404 by adding 10µl of plasmid to 50µl of *A. tumefaciens* competent cells. This solution was thoroughly mixed and submerged in liquid nitrogen. The tubes were heated to 37°C for five minutes. LB broth media (600µl) was added and it was then incubated for 2-3 hours at 28°C with shaking, then plated. After growing for 4 days at 28°C, a single colony was re-plated and grown for two days and used for both glycerol stocks and infiltration using the protocol used previously (Goodin et al., 2002). Time course experiments were conducted at two days to five days post-infiltration by were mounting leaf sections in water on a slide to prepare for visualization with the microscope,

### Expression of SVNV proteins in Sf9 insect cells

These SVNV pENTR-d-TOPO clones were also used for localization in insect cells. LR reactions with previously described and confirmed entry clones were conducted with pIWG (Teh et al., 2022). To identify SVNV proteins’ localization pattern, lepidopteran ‘SF9’ cells (Gibco™ Sf9 cells, Invitrogen) were transfected at approximately 85-90% confluency. Following the manfacturer’s recommendations, transfection was carried out with SVNV plasmids using Cellfectin II (Gibco™) in a 35 mm^2^ six well plate. Transfected cells were incubated at 28°C for 72 hours. A positive control of pHSP-70-GFP (Martin and Whitfield, 2018), as well as, a negative control (no plasmid) to determine the cell autofluorescence were both utlized. Each construct was localized between 3-5 times for microscopy, obtaining three photosets from each localization.

### Microscopy of plant and insect cells

To conduct these experiments, an Eclipse Ts2R (Nikon) epifluorescence microscope was used. Images were captured using the Eclipse Ts2R (Nikon) epifluorescence microscope and Nikon Elements software v. 5.21.02 (Build 1487). A final replicate was taken on the confocal microscope, Nikon A1R confocal with laser lines 405nm, 488nm, and 561nm, on a Nikon Ti-E inverted microscope stand using the C-Apochromat VC 40x/1.5 objective, for both the localization and co-localization in plant cells. The software on the confocal is Elements C, version number 4.40. For both the localization and co-localization experiments, image acquisition was conducted at 1024 × 1024 pixels, 12-bit depth, scan direction: one way, scanner zoom: 2.532-10.753, and scan speed: 0.125-0.25

Insect cell microscopy was carried out 72 hours after transfection using an Eclipse Ts2R (Nikon) epifluorescent microscope equipped with an S Plan Fluor ELWD 40x/.060 objective and a CFI 10x/22 eyepiece. 10 ng/µl DAPI stain solution resuspended in PBS pH 7 was utilized to stain the SF9 insect cells to visualize the nucleus, with staining occurring one hour prior to imaging. For both plant and insect cells, to ensure accurate representation, each of the images were meticulously compared during figure assembly, selecting the most appropriate one to depict the described results. Figures from both the Eclipse Ts2R (Nikon) epifluorescence microscope and the confocal microscope were compiled using Microsoft PowerPoint v. 2305 (Build 16501.20210 Click-to-Run).

### Field sample collection from Alabama

Soybean samples were collected from 33 fields in 14 counties in Alabama representing the northern and central parts of the state where the disease is most common (Figure 7-D). Counties that represent north Alabama include Blount, Cullman, Dekalb, Etowah, Jackson, Lawrence, Limestone, Madison, and Marshall. Counties that represent central Alabama include Chambers, Chilton, Elmore, Lee, and Talladega. Samples were collected preferentially based on vein necrosis symptoms suspected to be caused by SVNV regardless of the soybean cultivar. After collection, all samples were stored at -80°C until the total RNA was extracted.

The total RNA was extracted out of the 33 collected symptomatic soybean leaves. Taking into consideration that SVNV is not systemically moving in soybean by its own (Zhou and Tzanetakis 2020), the samples used for RNA extraction were preferentially collected from the necrotic tissues to guarantee SVNV is included. The symptomatic tissues (100 mg) were placed into 2 ml screw cap tubes containing 5-7 sterilized 2.4 mm metal beads. Tubes were then frozen in liquid nitrogen, and the frozen tissues were subsequently grounded using a Bead Ruptor Elite (Omni International, Kennesaw GA, USA). Grinding was carried out in two cycles, each lasting 30 seconds at 5 m/s, with the samples being re-frozen in liquid nitrogen between the cycles. RNA extraction was then conducted using the IBI Mini Total RNA Kit (Plants) (IBI Scientific, Dubuque, IA, USA) following the manufacturer’s protocol. The cDNA synthesis was performed using 1000 ng of total RNA and the Verso cDNA Synthesis Kit (Thermo Fisher Scientific, Waltham MA, USA) and the segment-degenerate reverse primer (Table 1) following the manufacturer’s instructions.

### Mechanical inoculation of SVNV

Wildtype *N. benthamiana* plants were grown and cultivated using the same method mentioned above (see above) and were used for mechanical inoculation of SVNV. Field collected, infected soybean samples from Alabama which tested positive for SVNV via PCR (described below) were used to conduct the mechanical transmission experiment. Soybean leaf tissues showing necrosis symptoms were ground in a sterilized mortar in 0.01 M sodium phosphate buffer, pH 7.0. *N. benthamiana* plants were dusted with Carborandum 600 mesh and the sap of the ground soybean tissue was then used for inoculation. At 14 days post-inoculation (dpi), inoculated and non-inoculated *N. benthamiana* leaf tissues were used for SVNV detection using the PCR.

### Soybean thrips collection and SVNV detection

In 2023, soybean thrips (*Neohydatothrips variabilis*) were collected in August 2023 from soybeans in the Cullers rotation managed by Auburn University in Lee County. Eighty insects were moved into a sterile 1.7mL tube containing 250µL of TRIzol (Ambion life technologies) and ground thoroughly. An additional 750µL of TRIzol was added while grinding and then incubated for three minutes using a rotating mixer. Samples were spun down at 12,000 x g at 4°C for 15 minutes. TRIzol was collected while carefully avoiding the insect debris and placed in a new sterile 1.7mL tube. Chloroform (200uL) was added and the tube was shaken vigorously for 15 seconds before incubating at room temperature for five minutes. Samples were then spun down at 12,000 x g at 4°C for 10 minutes. The aqueous phase (top layer) was carefully collected and placed in a new sterile 1.7 mL tube with 500µL of isopropanol. Samples were then shaken gently by hand and incubated at room temperature for 10 minutes. After incubation, the samples were spun down at 12,000 x g at 4°C for 10 minutes. The supernatant was removed, taking caution not to disturb the small white RNA pellet at the bottom. The pellet was washed with 500µL of 80% ethanol and spun down at 7,500 x g at 4°C for 5 minutes. All ethanol was removed and after two minutes at room temperature to dry the pellet was redissolved in 10μL of molecular grade water. Sample concentrations were checked and 1000ng of the total RNA was used to synthesize the cDNA using random hexamers and the Verso cDNA Synthesis Kit (Thermo Fisher Scientific, Waltham MA, USA) following the manufacturer’s instructions.

To amplify the three SVNV genes from thrips (N, NSs, and NSm), one microliter from the cDNA was used. Specific SVNV-N, NSs, and NSm PCR primers were used to amplify the three targeted genes (Table 1) using Phusion High-Fidelity PCR using the same methods as described above (see above). Purified PCR products were cloned into pJET-T-d-TOPO using the CloneJET PCR Cloning kit (Thermo Scientific Baltics UAB) following the manufacturer’s recommended instructions and confirmed with the pJET-F and pJET-R primers (Table 1) before sequencing. For some samples where pJET colonies could not be isolated, cloning was conducted with pENTR-d-TOPO using the methods described previously.

The sequences of SVNV three genes (N, NSS, and NSm) were analyzed using Blast (www.ncbi.nlm.nih.gov), by comparing SVNV-AL three gene sequences with SVNV gene sequences available on the NCBI. Sequences that have a full read with a start codon and no premature stop codons were only considered for the next steps. The sequences with full reads were translated from nucleotide sequence (DNA) to protein sequence (amino acids) using the Expasy tool, Translate (https://web.expasy.org/translate/). Samples of complete protein sequences with no gaps were then used to build the alignment using the Clustal Omega tool (https://www.ebi.ac.uk/Tools/msa/clustalo/) to detect the conserved and non-conserved amino acid mutations of SVNV of AL three gene with the Tennessee reference genome strain (GCF_004789395.1). The protein alignment was converted to a FASTA format using the online CONVERT tool (https://molbiol-tools.ca/Convert.htm).

### Construction of the phylogenetic trees

To indicate the likelihood between SVNV of Alabama and other SVNV sequences available on the NCBI, the phylogenetic tree was constructed using MEGA11: Molecular Evolutionary Genetics Analysis version 11 version 11.0.13. (Stecher et al., 2020). References available on NCBI including the TN genome for each gene were used to be compared to our sequences. Additionally, three important orthotospoviruses including Tomato spotted wilt virus (TSWV), Capsicum chlorosis virus (CaCV), and Bean necrotic mosaic virus (BeNMV) were also compared to our SVNV from Alabama on the phylogenic trees. This will allow us to understand the similarity of SVNV from Alabama to other members of the genus Ortotospovirus.

### SVNV population structure

The DNA and amino acid sequences of the nucleocapsid protein (NP) of the SVNV strains isolated from Alabama were used to calculate the identity percentage and the mean overall genetic distance. SVNV strains isolated from Alabama available on GenBank (accessions PP840132-63) were used along with 46 NP sequences from AR, DE, IL, KS, MD, MS, MO, KY, and TN (accessions HQ728355-84, HQ728386, JQ946869-74, JF808207-13, JF8082115, and JQ277450-52) collected between 2008 and 2012 (Zhou and Tzanetakis 2013). For the identity percentage, the DNA and amino acid sequences were aligned using the Clustal Omega tool (Madeira et al., 2024), and the percent identity matrix results were downloaded and viewed in Microsoft Excel (Version 16.88 “24081116”). The minimum and maximum identity percentages were then observed. For the mean overall genetic distance, the DNA and amino acid sequences were aligned using the ClustalW algorithm available in MEGA11: Molecular Evolutionary Genetics Analysis version 11.0.13. (Stecher et al., 2020). The alignments were then saved in meg format and analyzed for their overall mean distance.

## Results

### Domain analysis of SVNV genes

In the domain analysis of the nucleocapsid protein (N), the non-structural silencing suppressor protein (NSs), the non-structural movement protein (NSm), and the glycoproteins (G_N_/G_C_), of SVNV, we identified the presence of N-glycosylation sites across these viral proteins (Table 2). In SVNV, the glycoproteins G_N_ and G_C_ are hypothesized to be integral to viral entry and attachment to host cells (Whitfield et al., 2004). In SVNV, the cleavage between the polyprotein into separate glycoproteins, G_N_ and G_C,_ have been previously described as between Cys378 and Ser379 (Zhou and Tzanetakis, 2018).

**Table 2:** The size and predicted domains of Soybean vein necrosis virus proteins including the signal peptide (SP), transmembrane domain (TM), Nuclear Localization Signals (NLS), and N-Glycosylation sites.

| Protein | Size (aa) | Signal peptide (SP) (aa) | Transmembrane Domain (TM) (aa) | Nuclear Localization Signal (NLS) Sites (aa) | NLS Score | NLS type | N-Glycosylation Sites (aa) |
| --- | --- | --- | --- | --- | --- | --- | --- |
| N | 277 | None | None | 51-59<br>60-90<br>236-270 | 4<br>4.3<br>5 | Bipartite | 35-38, 218-221 |
| NSs | 440 | None | None | 333-363 | 4.8 | Bipartite | 158-161, 192-195, 288-291, 322-325 |
| NSm | 316 | None | None | 97-126 | 4.6 | Bipartite | 190-193 |
| G<sub>N</sub> | 451 | 6-28 | 324-343 | 104-134<br>416-447 | 4.5<br>4.7 | Bipartite | 25-28, 229-232, 343-346, 383-386, 398-401 |
| G<sub>C</sub> | 741 | 455-477 | 1105-1126 | None |  |  | 549-552, 1180-1183 |

Furthermore, three bipartite Nuclear Localization Signals (NLS) were identified in the nucleocapsid protein with scores 4, 4.3, and 5, One bipartite NLS was identified in the NSs protein with a score of 4.8, One bipartite NLS was identified in NSm with a score of 4.6 and two bipartite NLS were identified in G_N_ with a score of 4.5 and 4.7 (Table 2). G_C_ had five NLSs with scores ranging from 3 to 3.8, but we determined our threshold for NLS scores was below four as these were less likely to be relevant. Additionally, a signal peptide domain (SP) and a transmembrane domain (TM) were detected in both G_N_ and G_C_ (Table 2). These domains will likely target these proteins to the membranes of the cell which is expected when compared to other glycoproteins in orthotospoviruses (Dietzgen et al., 2012, Tripathi et al., 2015, and Martin et al., 2024).

### Localization of SVNV ORFs in N. benthamiana

To confirm the domain analysis and further understand this virus, subcellular localization of SVNV proteins was done. We were expecting some differentiations for this virus’s protein localizations as it is present in a unique clade with BeNMV separate from other othotospoviruses (Whitfield 2016 and Kormelink et al., 2021). We expected to find SVNV-N protein localizing to the cell periphery consistent with the cytoplasm. As N has RNA-binding domains, cytoplasmic localization is needed to encapsulate the viral RNA during morphogenesis (Li et al., 2014). Interestingly, GFP-N localized to the cell periphery and the plant cell nucleus (Figure 2: I-III). RFP-N is localized to the cell periphery but not to the cell nucleus (Data not shown), however, this may be due to a lesser fluorescence intensity in the nucleus with RFP compared to GFP (Martin et al., 2009). N-GFP and N-RFP localized to the cell periphery but was weekly localized in/around the cell nucleus (Supplementary Figure 1-A, I-III for N-GFP, and Supplementary Figure 2-B, I-XV for N-RFP). Although NLS signals were predicted for N, the scores were low, thus this localization was somewhat unexpected.

**Figure 2:**
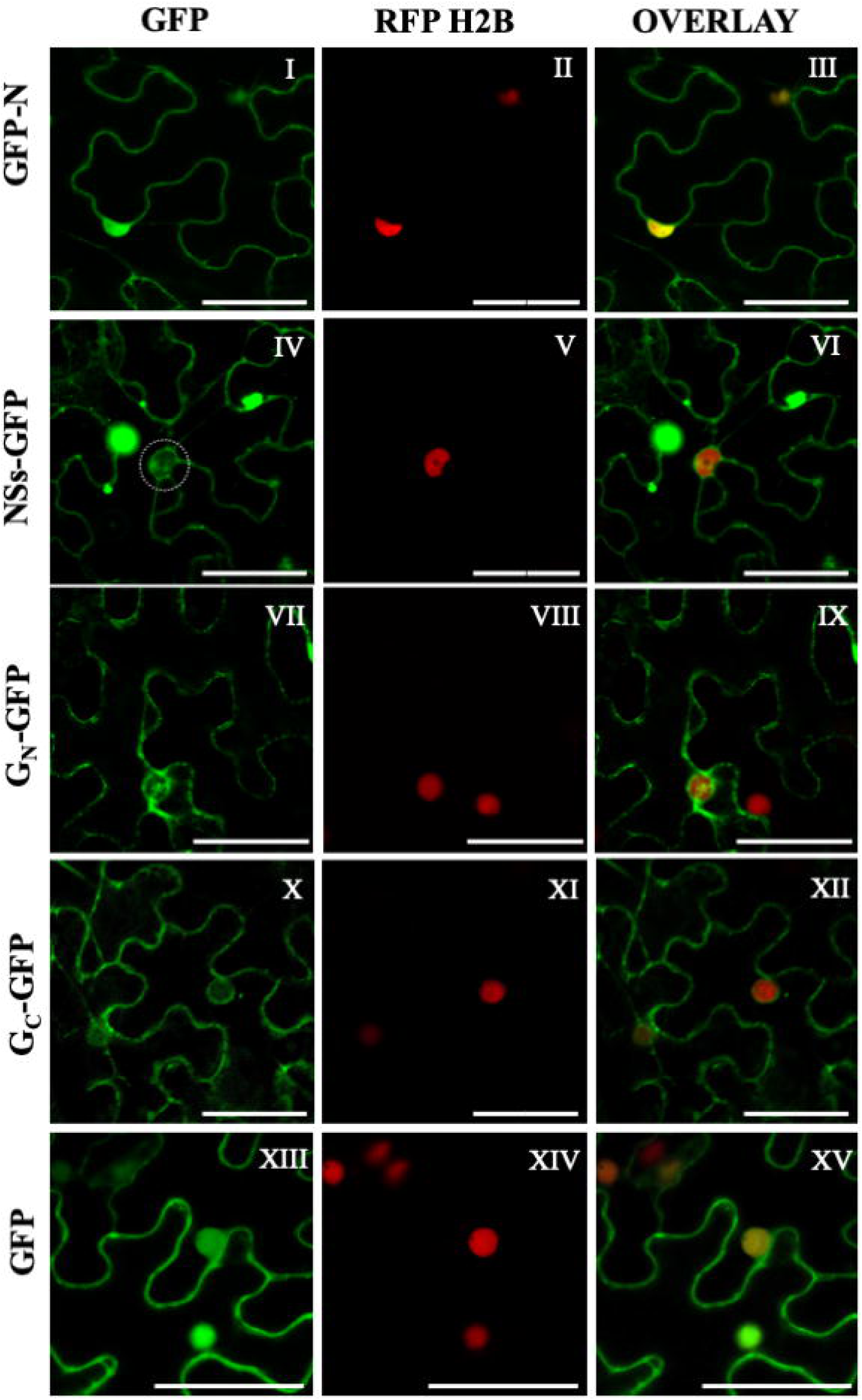
The localization of the Soybean vein necrosis virus proteins (SVNV) in plant cells. from left to right, column 1, is the GFP localization for each of SVNV proteins, column 2, is the cellular marker RFP-H2B, and column 3, is an overlay of columns 1 and 2. The rows from left to right are: I-III, GFP-N, IV-VI, NSs-GFP, VII-IX, GN-GFP, X-XII, GC-GFP, and XIII-XV, the free-GFP localization control. The scale bar is 50µm.

For the non-structural silencing suppressor (NSs), we expected to see cytoplasmic/cell periphery localization based on previously localized othotospoviruses (Dietzgen et al., 2012, Tripathi et al., 2015, and Martin et al., 2024). In our experiments, SVNV-NSs localized to the cell periphery and aggregates in the cytoplasm when fused to the C terminus of GFP and RFP (Data not shown). NSs-GFP localized to the cell periphery and weakly in the cell nucleus which was not expected (Figure 2: IV-VI). However, this does correlate to the presence of NLSs signals which were predicted for this protein, although with a median score of 4.8 out of 10 indicating both cytoplasmic and nuclear localization (Kosugi et al., 2008, Kosugi et al., 2009a, and Kosugi et al., 2009b).

Based on the comparison of domains of SVNV to TSWV, we expected to observe the localization of SVNV-NSm in the plasmodesmata (PD). Some variation was expected as SVNV does not systemically move in soybean on its own but does in the presence of other soybean viruses (Zhou and Tzanetakis 2020). GFP-NSm, RFP-NSm, NSm-GFP, and NSm-RFP did not show protein expression after eight different localization replicates. After 5dpi, necrotic symptoms starting as yellowing on the infiltrated leaves and ending in brown necrotic spots were seen (Data not shown). Suspecting cell death, we utilized SYTOX blue stain, which determines plant cell viability in living plant samples using fluorescence microscopy. SYTOX blue stain and its spectral variants have the ability to bind to DNA but cannot penetrate the plasma membrane of viable plant cells (Truernit and Haseloff 2008). Infiltrated cells with NSm-GFP showed the SYTOX blue was inside the cells indicating cell death (Figure 3: I-IV). GFP-N was used as a control and the SYTOX was not present in the cells, but outside (Figure 3 V-VIII). This indicates that the cell death observed during infiltration/infection may be due to NSm.

**Figure 3:**
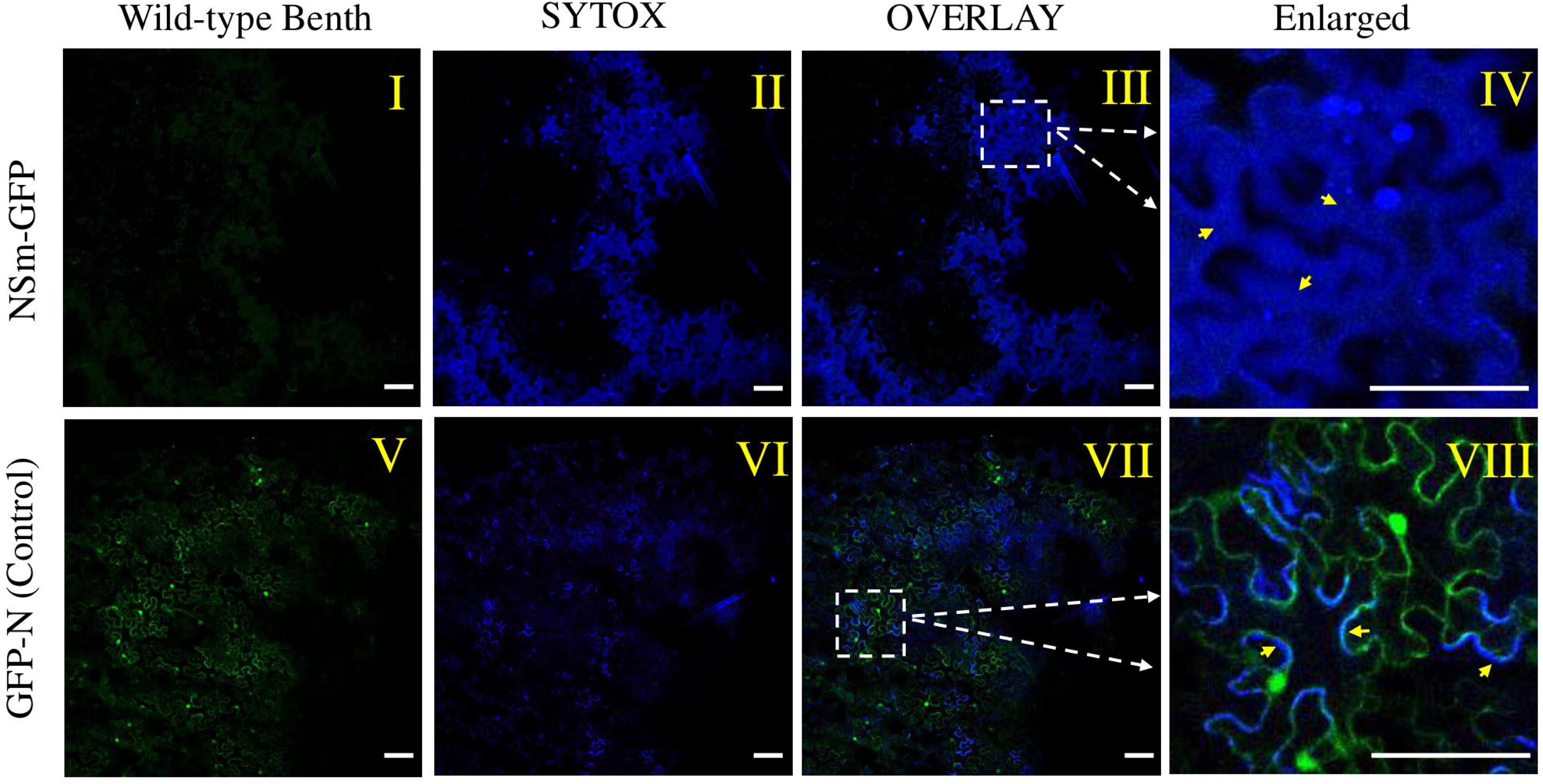
The confocal microscopy images of the SYTOX blue stain used to test plant cell viability taken at 3dpi after the Nicotiana benthamiana leaves were infiltrated with SVNV-NSm-GFP. Columns from right to left are the first column is the wild-type of N. benthamiana, the second column is SYTOX/1000nMOL blue stain infiltrated 30 minutes in the leaves before imaging, the third column is an overlay with columns one and two, and the fourth column is an enlarged section of images highlighted with dotted boxes. Rows from top to bottom are, I-IV, NSm-GFP, V-VIII, GFP-N used as a control. The scale bar is 100µm

To avoid blocking the signal peptide function in the glycoproteins, G_N_ and G_C_ (Figure 1), they were fused exclusively to the N terminus of GFP and RFP. Both G_N_ and G_C_ contain a transmembrane domain embedding them in the cell membrane, thus we expected to see both proteins in the outer and internal membranes. Both SVNV-G_N_-GFP and G_C_-GFP are localized to the cell membrane and around the cell nucleus (Figure 2: VII-IX, X-XII), which agreed with our hypothesis. The Free-GFP was used as a localization control, and it localized to the cell periphery and inside the nucleus as expected (Figure 2: XIII-XV).

### Co-Localization of SVNV ORFs in N. benthamiana

*N. benthamiana (*wild-type) plants were used to image two SVNV proteins infiltrated together to test the impact of the presence of one protein with other proteins. All co-localizations were done for SVNV proteins using the N terminal fusion to GFP and RFP. At 2 dpi, when NSs-GFP was co-localized with N-RFP, both proteins co-localized to the cell periphery and the cell nucleus (Figure 4: I-III), and this changed after 3 dpi, and therefore, a time course was taken recording this change until 6dpi. NSs-GFP and N-RFP were localized alone, and same time course images were taken until 6 dpi to compare it with the co-localization time course. For NSs-GFP, it localized to the cell periphery and weakly to the cell nucleus from 2 – 4 dpi (Supplementary Figure 2-A: I-IX). At 5 – 6 dpi, very bright aggregates of NSs were seen outside the nucleus (Supplementary Figure 2-A: X-XV). For N-RFP, from 2 – 6 dpi, it localized to the cell periphery and weakly to the cell nucleus in every instance (Supplementary Figure 2-B: I-XV). When both proteins are infiltrated together, they co-localized to the cell periphery and the cell nucleus at 2 dpi. Additionally, an area with no protein localization formed (voids) by both NSs and N around the nucleus (Supplementary Figure 2-C: I-IV). At 3dpi, NSs started forming small aggregates two days earlier than when it was alone (Supplementary Figure 2-C: V-VIII). From 4 dpi to 6 dpi, both proteins accumulated around the nucleus after 3 dpi to 6 dpi (Supplementary Figure 2-C: IX-XX). N-RFP which always localized to the cell periphery and weakly to the cell nucleus, showed protein aggregations similar in size to those formed by NSs-RFP when it was present with NSs-GFP, which has never seen with N-RFP alone.

**Figure 4:**
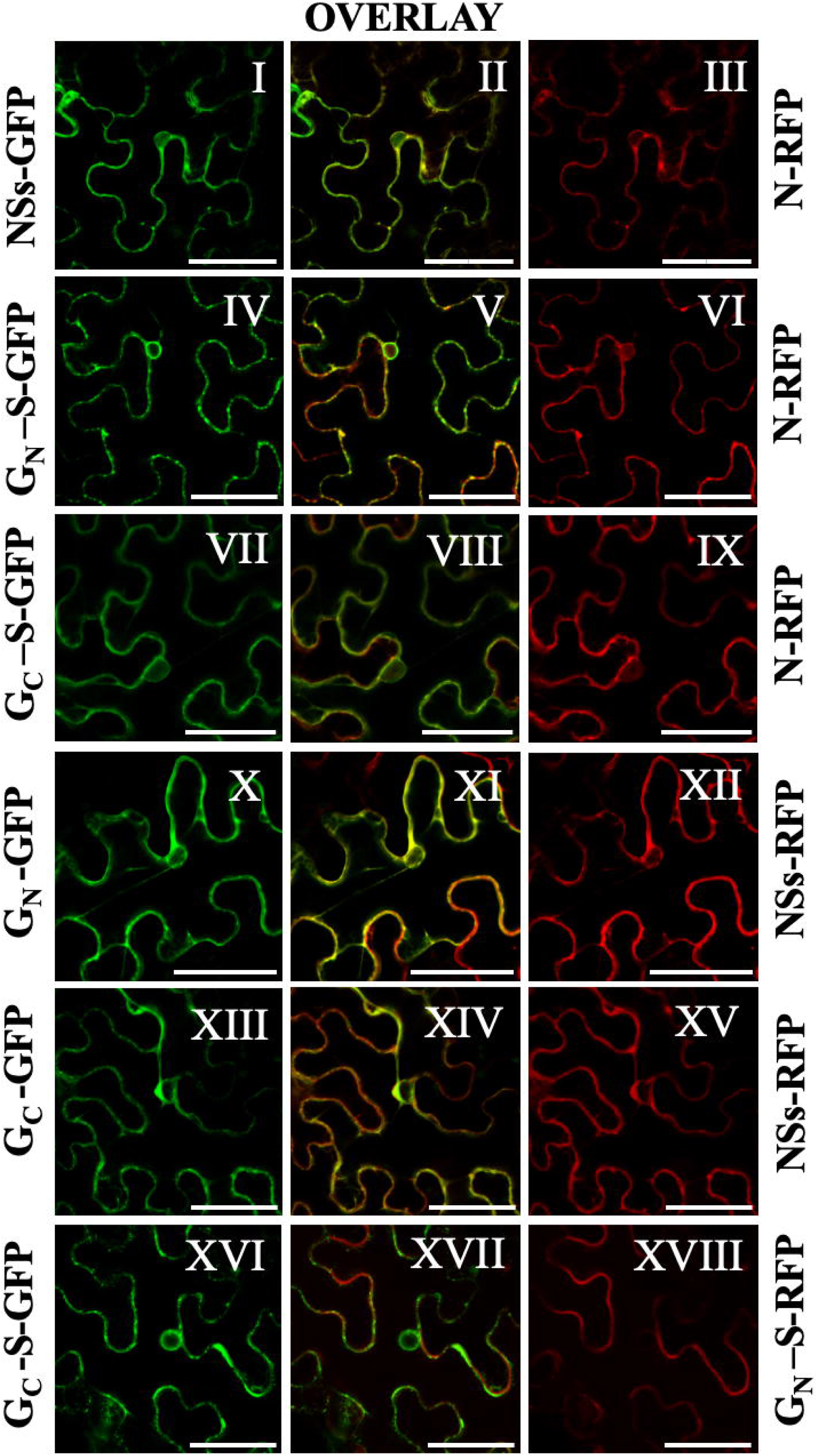
The co-localization of the Soybean vein necrosis virus proteins (SVNV) in plant cells at 2 dpi. From left to right, column 1, is the SVNV protein tagged with GFP, column 2, is an overlay with columns 1 and 3, and column 3, is the SVNV protein tagged with RFP. The rows from left to right are, I-III, NSs-GFP with N-RFP, IV-VI, GN-S-GFP with N-RFP, VII-IX, GC-S-GFP with N-RFP, X-XII, GN-GFP with NSs-RFP, XIII-XV, GC-GFP with NSs-RFP, and XVI-XVII, GC-S-GFP with GN-S-RFP. The scale bar is 50µm.

When NSs-RFP was co-localized with G_N_-GFP and G_C_-GFP, both G_N_-GFP and G_C_-GFP could be more easily visualized than with N-RFP (Figure 4: X, XIII, IV, and VII). There was no change between the co-localization of G_N_-GFP and G_C_-GFP in the presence of NSs-RFP and the individual localization for each of them as they localized to the cell periphery/membrane and around the cell nucleus in both experiments (Figures 2: VII-IX and X-XII and Figure 4: X-XII and XIII-XV). The protein expression of the full-length forms of both G_N_ and G_C_ was very weak, thus the soluble forms were used. G_N_-S-RFP was co-localized with G_C_-S-GFP, as these are expected to interact during infection and both co-localized to the cell membrane/periphery. Additionally, G_C_-S-GFP localized around the cell nucleus (Figure 4: XVI-XVIII).

When NSm was localized alone, it did not express (Figure 3: I) and caused cell death (Figure 3: I-IV). To test if the presence of the other viral proteins results in stabilizing the cell environment for the expression of NSm (Chen and Zacharias 2023), we conducted co-localization experiments. Following this hypothesis, we co-infiltrated the NSm with SVNV proteins (N, NSs, G_N_, G_N_-S, G_C_, and G_C_-S), however, NSm still did not express, and the other proteins did not change their localization patterns compared to when they were alone. Interestingly, when NSm-RFP was co-infiltrated with G_N_-GFP and G_C_-GFP, there was a greater expression and also aggregations seen at two dpi for both G_N_-GFP and G_C_-GFP, with no expression for NSm-RFP (Data not shown). This glycoprotein over-expression and aggregation were not recorded before either when they were localized individually (Figure 2: VII and X) nor when they were co-localized with N-RFP or NSs-RFP (Figure 4: IV, VII, X, and XIII). This experiment was repeated three times, and the same results were recorded in every replicate. It is unclear if this response is due to the presence of NSm in some small quantity or the possibility that cell death itself caused this response.

### Localization of SVNV in insect cells

To better understand SVNV’s ability to infect both plant and insect hosts, we investigated the subcellular localization of SVNV proteins within insect cells. For this study, Sf-9 cells were utilized for the cellular environment of thrips, the natural insect host for SVNV, due to the unavailability of thrips cell lines. The nucleocapsid (N) protein is predominantly localized to the cytoplasm, particularly around the nucleus (Figure 5: I-V). This localization was expected, as N has been shown to contain an RNA binding domain (Li et al., 2014) and N would need to be present in the cytoplasm to encapsulate genomic RNA. Comparatively, in *N. benthamiana* plant cells, the N showed localization in the cell periphery and in the cell nucleus (Figure 2: I-III), indicating a slightly different localization. It should be noted that the nuclear localization seen in plants was in an orientation not tested in insect cells and that the N-GFP in plants also did not localize to the nucleus but around it (Supplementary Figure 1: I-III).

**Figure 5:**
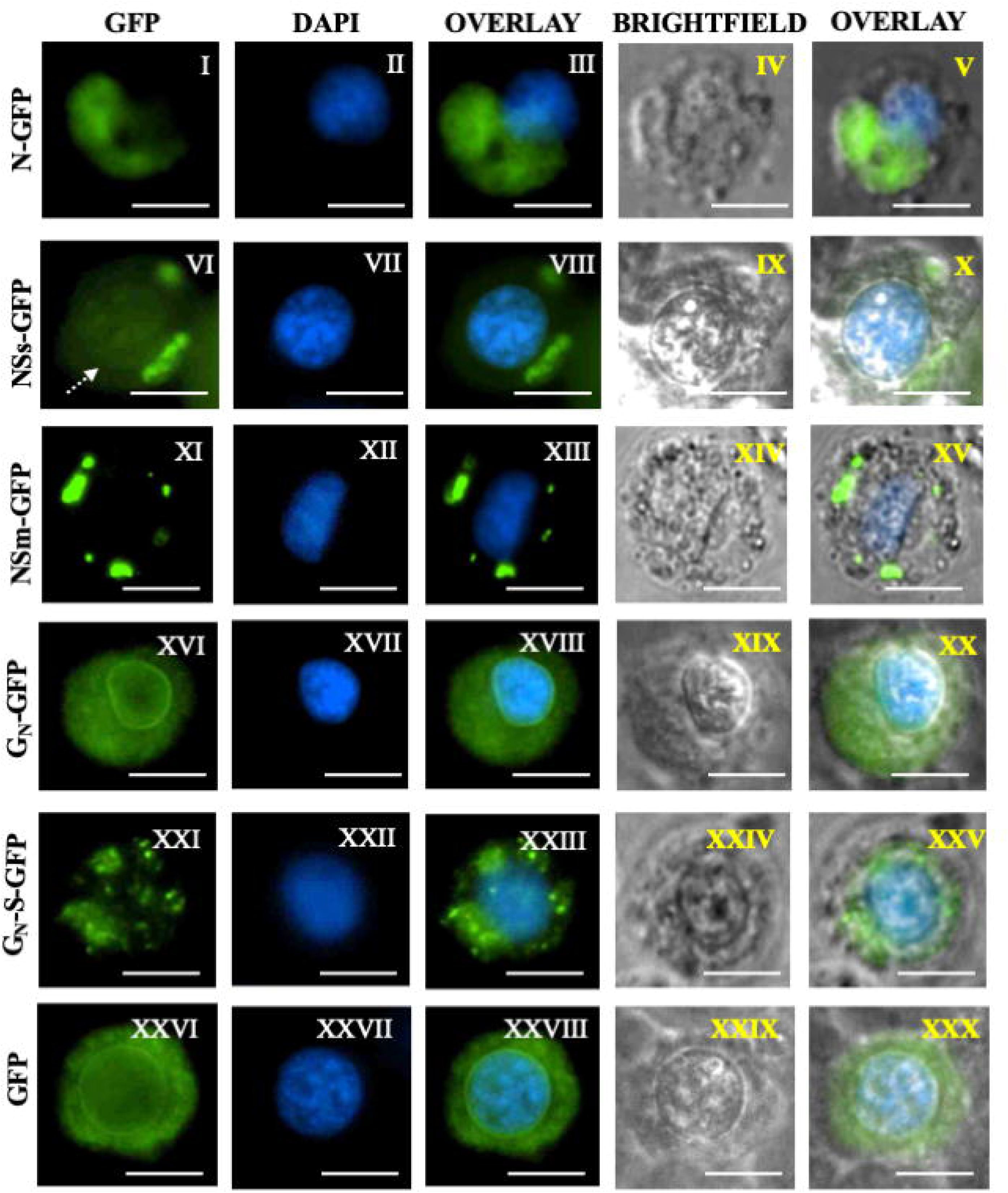
Localization of SVNV proteins in SF9 insect cells. Column 1, GFP tagged to SVNV protein, Column 2, DAPI stain to indicate nucleus, Column 3, overlay of Columns 1 and 2, Column 4, brightfield, Column 5, overlay of Columns 1-4. Rows are as follows: I-V, SVNV-N-GFP, VI-X, SVNV-NSs-GFP, XI-XV, SVNV-NSs-GFP, XVI-XX, SVNV-GN-GFP, XXI-XXV, SVNV-GN-S-GFP, XXVI-XXX, pSHP70 Free-GFP control. The scale bar is 25µm.

NSs also exhibited cytoplasmic localization (Figure 5: VI-X). Unlike N, NSs showed aggregations (Figure 5: VI-X) that ranged in size similar to plant cells (Figure 2 IV-VI). Similar to the localization in plant cells (Figure 2: IV-VI), NSs was seen to weakly localize in the cell nucleus of the SF9 insect cells. It is expected that NSs is not retained in the nucleus due to the weak localization (Figure 5: VI-X). In the *in silico* NLS signal predictor, there was a single NLS site found (Table 2), indicating that there is a possibility for nuclear localization in insects as well as plants.

Interestingly, the non-structural movement protein (NSm) displayed cytoplasmic localization (Figure 5: XI-XV) but appeared less uniform than the N or NSs, forming aggregates within the cytoplasm (Figure 5: XI-XV). While the localization pattern was consistent across all replications. In contrast, in *N. benthamiana* plant cells, NSm could not be localized and instead caused cell death in the infiltrated cells (Figure 3: I-IV).

Similar to the localization in plant cells (Figure 2: VII-IX), the glycoprotein G_N_ was dispersed throughout the cytoplasm (Figure 5: XVI-XX) and prominently surrounding the nucleus, localizing to the membrane around the nucleus of the cell (Figure 5: XVI). This localization and membrane association was expected, as full-length G_N_ contains a transmembrane domain. G_N_-S localized in the cytoplasm (Figure 5: XXI-XXV) and formed small aggregates (Figure 5: XXI) consistent with its lack of transmembrane domain (Table 2). It is also noticeable that G_N_ form aggregates, but rather localizes all throughout the cytoplasm. Surprisingly, we were unable to localize G_C_, even when the transmembrane domain was deleted (G_C_-S). Out of 10 replicates, both G_C_ and G_C_-S, localization was not seen.

### Soybean sample collection, gene amplification and cloning, and Sanger sequence analysis

In 2023, a total of 33 soybean samples were collected from SVNV symptomatic soybean fields in 14 different counties in AL. All 33 samples tested positive for the three targeted genes of SVNV. The soybean thrips collected from one location also tested positive for N, NSs, and NSm genes. For the N gene, 31 complete sequences were obtained, however for NSs gene, 25 complete sequences were obtained. For the NSm gene, 30 complete sequences were obtained. Moreover, a complete gene sequence of N and NSm was obtained from the soybean thrips, however, the cloning of the NSs gene from thrips was not successful. A complete sequence is defined as a sequence which contains a full ORF with no premature stop codons interrupting the sequence that matches to SVNV on the NCBI database. All partial sequences and those not matching to SVNV were not considered.

### Mechanical inoculation of SVNV

In an interest of exploring the systemic spread of SVNV, we mechanically inoculated *N. benthamiana* plants with field collected samples. We expected to detect SVNV in locally inoculated *Nicotiana benthamiana* leaves but not in the top or lower non-inoculated leaves because SVNV was reported that it does not move systemically on its own (Zhou and Tzanetakis 2020). *N. benthamiana* mechanically inoculated plants with SVNV were tested using the PCR with the same PCR settings as mentioned before (see above). SVNV-N gene was detected from inoculated and non-inoculated *N. benthamiana* leaf tissues (Data not shown). This suggests that NSm from field samples is potentially marginally functional in *N. benthamiana* or not needed for systemic movement in that system.

### Protein alignment of SVNV genes

The protein alignment of the sequenced three genes of SVNV was conducted to determine sequence changes compared to earlier samples deposited in NCBI. We expected mutations in the virus genome when it is compared to its reference genome simply due to the length of time since the original sequences were deposited. The protein alignment of SVNV-N indicated the presence of eight conserved and non-conserved amino acid mutations compared to the reference genome of TN (GCF_004789395.1) (Figure 6-A and Supplementary Table 1). Of the eight mutations: position 13, Alanine (A) changed to Valine (V)(30/32, 93.75%); position 15, Isoleucine (I) changed to Leucine (L)(32/32, 100%); position 22, Lysine (K) changed to Arginine (R)(29/32, 90.62%); at Position 115, Alanine (A) changed to Glutamate (E) (28/32, 87.5%); position 118, Aspartate (D) mutated to Glutamate (E) (19/32, 59.37%); position 128, Leucine (L) changed to Phenylalanine (F)(18/32, 56.25%); position 169, Glycine (G) changed to Asparagine (N) (27/32, 84.37%); and lastly at position 227, Isoleucine (I) changed to Methionine (M) (22/32, 68.75%). The same eight amino acid mutations were detected in N of SVNV sequenced from soybean thrips (Supplementary Table 1).

**Figure 6:**
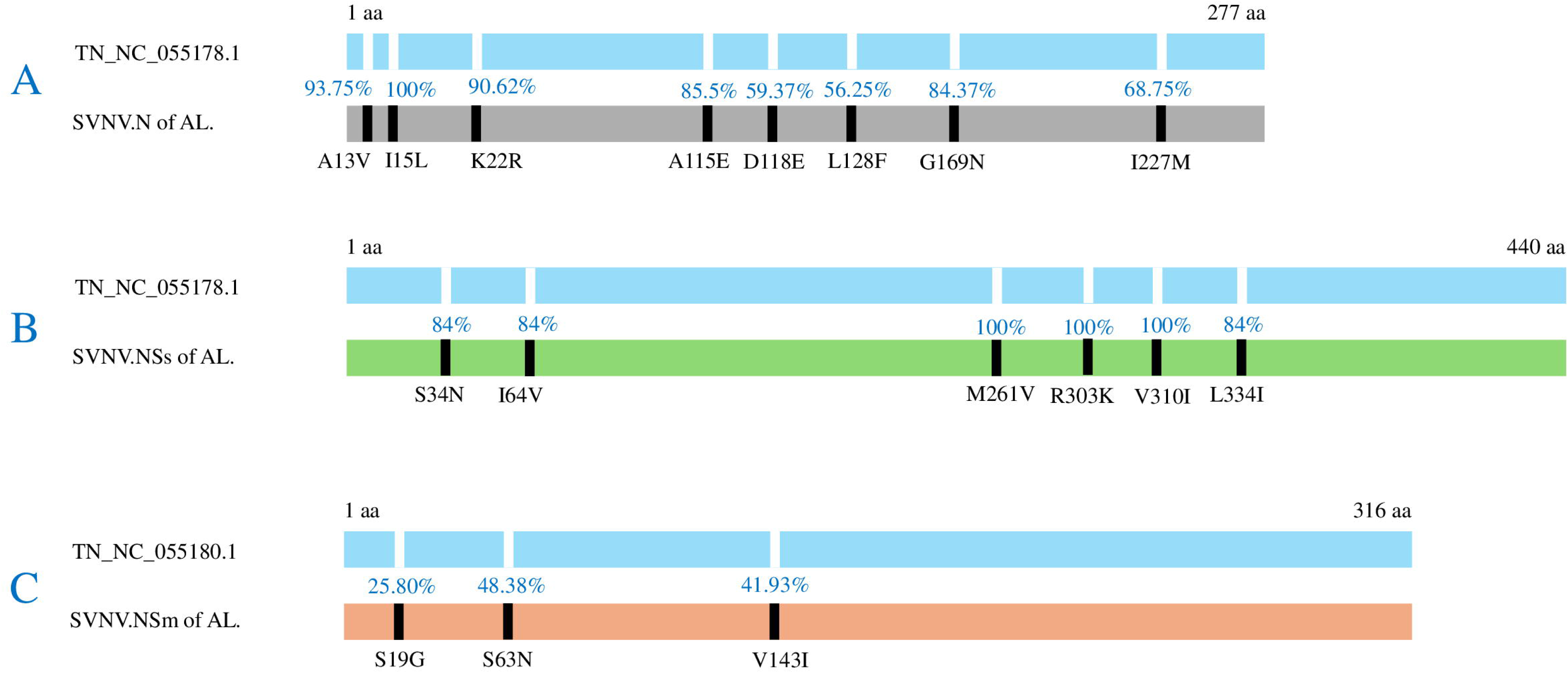
Conserved amino acid mutations that were observed in SVNV proteins collected from infected soybean fields in Alabama and the sites for each mutation; A: shows SVNV.N protein, B: shows SVNV.NSs protein, and C: shows SVNV.NSm protein. In each A, B, and C the top map is the comparison between the NCBI reference genome of the Tennessee strain and our strains of AL. The percentages are the numbers of our samples that show each of the mutations compared to the NCBI reference genome of the Tennessee strain.

**Figure 7:**
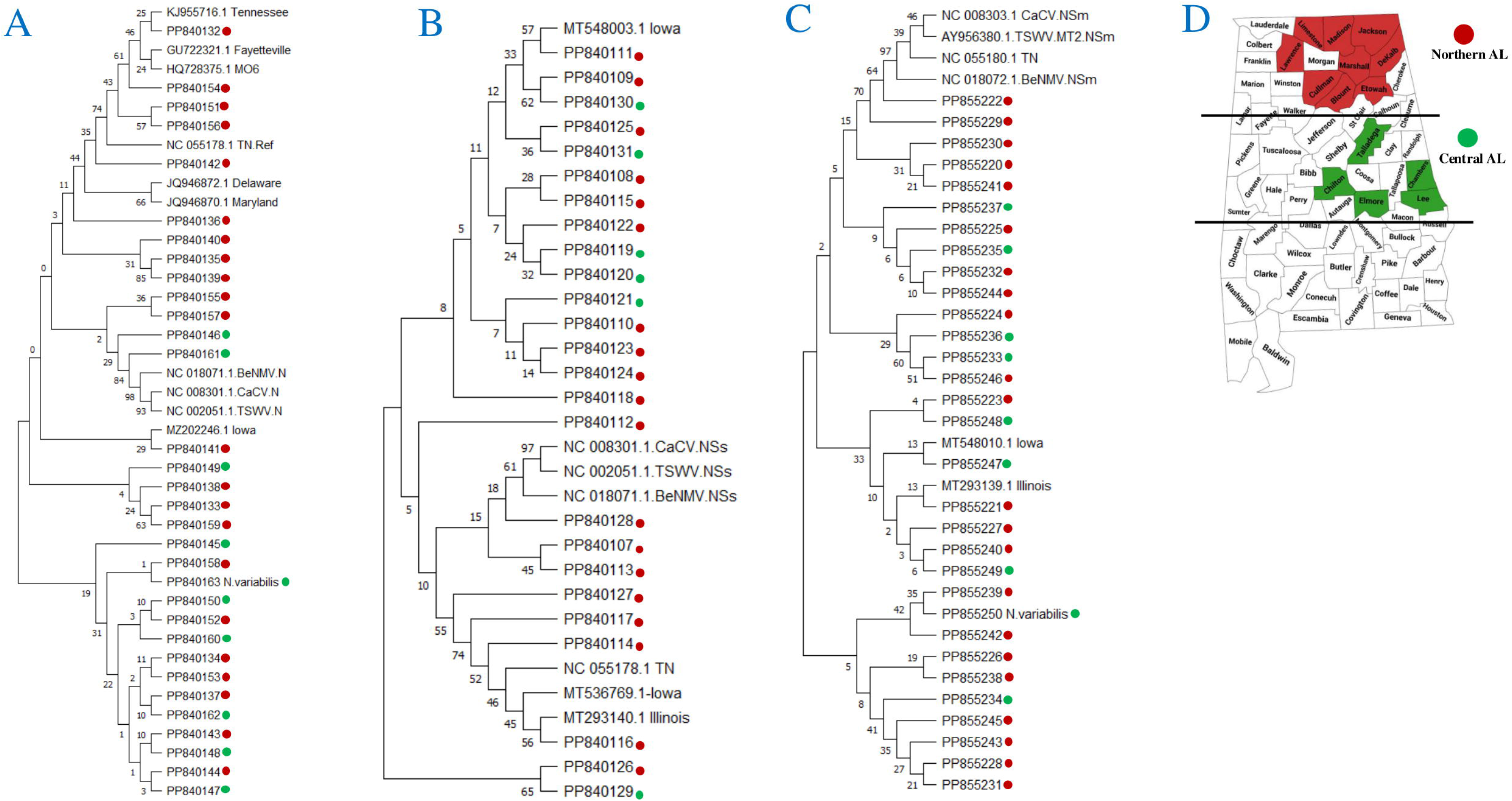
Phylogenetic tree of SVNV proteins collected from infected soybean plants in Alabama with the NCBI references including the reference genome of Tennessee. Bean necrotic mosaic virus (BeNMV), Tomato spotted wilt virus (TSWV), and Capsicum chlorosis virus (CaCV) were used as out-grouping orthotospoviruses. A represents the phylogenetic tree of SVNV.N protein, B represents SVNV.NSs protein, C represents SVNV.NSm protein, and D is a map of AL showing the counties used for samples collection from the north of AL marked in red color, and the counties used for samples collection from the center of AL marked in green color.

The protein alignment of SVNV-NSs revealed six conserved amino acid mutations that compared to the reference genome of TN (GCF_004789395.1) (Figure 6-B and Supplementary Table 1). At position 34, Serine (S) mutated to Asparagine (N) (21/25, 84%). Isoleucine (I) mutated to Valine (V) at site 64 (21/25, 84%). Methionine (M) mutated to Valine (V) at site 261 (25/25, 100%). Arginine (R) mutated to Lysine (K) at site 303 (25/25, 100%). Valine (V) mutated to Isoleucine (I) at site 310 (25/25, 100%). At site 334, Leucine (L) mutated to Isoleucine (I)(21/25 84%) (Figure 6-B and Supplementary Table 1).

The protein alignment of SVNV-NSm showed three partially conserved amino acid mutations compared to the reference genome of TN TN (GCF_004789395.1) (Figure 6-C and Supplementary Table 1). At site 19, Serine (S) changed to Glycine (G) (8/31, 25.80%) at position 63, Serine (S) is now to Asparagine (N) (15/31, 48.38%). Lastly, at site 143, Valine (V) changed to Isoleucine (I) (13/31, 41.93%). The first mutation was not detected in NSm identified from the soybean thrips, however, the second and third mutations were detected (Supplementary Table 1).

### Phylogenetic analysis of SVNV three genes

SVNV-N sequences that represent northern AL were phylogenetically closest to each other. The same trend was seen in sequences that came from central AL. However, not all northern or central samples followed this pattern, as some samples from both locations were also seen in mixed clades (Figure 7-A). Furthermore, the NCBI reference including Arkansas, Delaware, and Maryland were closest to SVNV from northern AL. The three out grouping orthotospoviruses were found in a clade that has both SVNV sequences from the northern and central of AL. Interestingly, N that came from soybean thrips from central AL (Lee County) was found closely related to sample N30 (PP840158) which came from the north of AL. (Figure 7-A).

Compared to the SVNV-N phylogenetic tree, the NSs phylogeny tree has less segregation (Figure 7-B). The TN reference genome (GCF_004789395.1) and the NCBI references including Iowa and Illinois were phylogenetically closest to the SVNV of northern AL. The three out grouping orthotospoviruses found closest to NSs sequences that came from northern AL. (Figure 7-B). Sequences from both sides of AL were more likely to be mixed rather than grouped based on northern or central sequences in the phylogenetic tree from NSm (Figure 7-C). In contradiction with the other two genes N and NSs, the TN reference NSm was found in a mixed clad with both samples from the northern and central AL. Moreover, the reference sequences from Iowa and Illinois were also in this mixed clade. Following the same trend as NSs, the three out-group orthotospoviruses were found closest to SVNV sequences that came from northern AL. (Figure 7-C).

### SVNV population structure

We further analyzed the sequences to determine if SVNV strains currently are similar to those collected previously and were homogeneous (Zhou and Tzanetakis, 2013) or if diversity increased over time. To answer this question, the DNA and protein sequences of our strains were added to the previous 46 SVNV sequences, and we found that the DNA sequences of all isolates of the SVNV-N gene show identities of 90.97% - 100%. On the other hand, the protein sequences of all isolates of SVNV-N show identities of 81.23% - 100%. The mean overall genetic distance between all isolates based on the DNA alignment was 0.01, and 0.02 based on the DNA alignment. This shows that the virus diversity and the genetic distance increased when we added our stains to the previous comparison (Zhou and Tzanetakis 2013).

## Discussion

Initially, we hypothesized that the localization pattern of SVNV proteins is likely to be similar, but not identical, to other characterized orthotospoviruses. To do this, we conducted the first localization of the ORFs N, NSs, NSm, G_N_ and G_C_. L was not included due to the size (9,010 bp). The GFP-N and N-RFP showed nuclear localization which has never been shown before in any localized orthotospovirus (Dietzgen et al., 2012, Tripathi et al., 2015, Widana Gamage and Dietzgen 2017, and Martin et al., 2024). N of the four characterized orthotospoviruses was localized to the cell periphery and aggregates in the cytoplasm. Additionally, in IYSV and CaCV,N was localized around the nucleus (Tripathi et al., 2015 and Widana Gamage and Dietzgen 2017). In support of this localization, N has three predicted bipartite NLS signals, one of them with a score of five indicating localization in both the cytoplasm and nucleus but not strong nuclear localization with the highest possible score of ten (Kosugi et al., 2008, Kosugi et al., 2009a, and Kosugi et al., 2009b) (Table 2). These nuclear domains may be implicated in the nuclear localization of SVNV-N in only one orientation (Supplementary Figure 1-A, I-III). The highest NLS domain score from N spanned from 236-270 aa (Supplementary Figure1-B), which is seven aa away (this includes the linker) from the start of GFP. This NLS domain could be blocked by GFP folding resulting in more cytoplasmic localization as observed in N-GFP (Supplementary Figure 1-A, I-III) and N-RFP (Supplementary Figure 2-B, I-XV). SVNV is the only orthotospovirus that has shown N nuclear localization, and it is unclear if the nuclear localization of SVNV-N plays a role in the virus replication or infection cycle or is interacting with host factors which cause movement to the nucleus or both.

The domains of SVNV-N of TN such as an RNA-binding Lysine-rich motif KKDGKGKKSK at positions 264–273, and several RNA-interacting amino acids (PSN7–9, RK51–52, RY54–55, and KK73–74) (Zhou and Tzanetakis 2019) may allow the nucleoprotein assist in RNA replication with the RdRp as shown for members of the *Bunyavirales* (Dunn et al., 1995 and Kainz et al., 2004). By looking into our eight conserved amino acid mutations that were observed in SVNV-N of AL, they do not fall at any of the reported motifs and the reported RNA binding regions. This might suggest that these motifs are crucial for the protein function and cannot change. Since three of the mutations we did find were conserved in all of our N sequences (100%) and three were 84% (Figure 6-A), they might enhance the self-interaction of the SVNV-N as previously been reported for TSWV, CaCV-AIT (Thailand), and INSV (Lacorte et al., 2007, Zilian and Maiss 2011, Dietzgen et al., 2012) additionally, for BeNMV, Chrysanthemum stem necrosis virus (CSNV), Iris yellow spot virus (IYSV), and Tomato chlorotic spot virus (TCSV) (Leastro et al., 2015; Tripathi et al., 2015).

The RNA silencing suppressor activity of NSs in plants has been shown, while its role in animal-infecting bunyaviruses remains a subject of debate (Hedil and Kormelink, 2016). NSs of INSV and IYSV was localized to the cell periphery. For TSWV-NSs, it was localized around the nucleus as well as to the cell periphery (Martin et al., 2024). The first documentation of the nuclear localization of the NSs among the four characterized orthotospoviruses was in CaCV (Widana Gamage and Dietzgen 2017). Our SVNV-NSs, in agreement with CaCV-NSs, is located in the cell periphery in plant cells and weakly in the cell nucleus in plant cells. It was also found in the nucleus of insect cells; however, this has only been tested for TSWV where NSs is cytoplasmic (Martin et al., 2024). The NLS Mapper detected one nuclear domain in the SVNV-NSs, (Table 2) and the nuclear localization of the NSs was not as bright as it was seen with N. We are confident it is nuclear as the nucleolus was seen as a dark hole in the GFP. This indicates NSs is not present in the nucleolus but inside the nucleus (Figure 2: IV). This weak nuclear localization was also seen in insect cells (Figure 5: VI). The weakness of the expression indicated NSs may not be retained in the nucleus. SVNV-NSs may shuttle between the cell nucleus and cytoplasm by interacting with unknown host factors/proteins (Widana Gamage and Dietzgen 2017), similar to the mechanism that tombusvirus P19 partially translocates to the nucleus using ALY proteins (Canto et al., 2006). However, presence or absence of the NLS domains in NSs may not be a strong indicator of nuclear localization, as TSWV-NSs has a predicted NLS domain but exclusively localized to the cytoplasm (Kormelink et al., 1991; Kikkert et al., 1997). The possible nuclear localization of SVNV-NSs needs to be validated using similar immunofluorescent techniques that were previously used by Widana Gamage and Dietzgen 2017.

The SVNV-NSs from TN contains conserved GK_178–179_ and DE_148–149_ which comprise the Walker A and B motifs (Zhou and Tzanetakis 2019). These important motifs interact with ATP/ADP phosphates and coordinate/bind Mg2^+ions^ during the ATP hydrolysis process (Caruthers and McKay 2002 and Lokesh et al., 2010). However, the six amino acid mutations that were detected in our SVNV-NSs of AL did not appear in these motifs and that the Walker A and B motifs remained conserved in our sequences. This might indicate the importance of these motifs for the functionality of the NSs. However, the detected six amino acid mutations in our sequences were conserved among our sequenced samples (Figure 6-B). At this time, the six conserved mutations require further exploration to determine their significance or if they are simply a marker for the amount of time that has passed since sequencing.

NSm of four orthotospoviruses (INSV, IYSV, CaCV, and TSWV) showed localization in the plant plasmodesmata (PD) which helps the virus to move between cells (Dietzgen et al., 2012, Tripathi et al., 2015, Widana Gamage and Dietzgen 2017, and Martin et al., 2024). The NSm of IYSV was also localized around the cell nucleus (Tripathi et al., 2015). Unique to SVNV, NSm (in both GFP and RFP and both orientations) did not show protein expression after more than eight separate infiltration trials. SVNV-NSm caused cell death on infiltrated cells, as shown by the SYTOX stain (Figure 3: I-IV). The SYTOX blue stain was seen inside the cells infiltrated by NSm-GFP indicating cell death, however, outside the cells with GFP-N indicating viability (Truernit and Haseloff 2008). The phylogenetic trees constructed by de Oliveira et al., 2012, Oliver and Whitfield 2016, and Kormelink et al., 2021) have confirmed that SVNV is phylogenetically closest to BeNMV and distinctly different from other orthotospoviruses. The placement of SVNV with BeNMV suggests that SVNV is part of a novel evolutionary lineage of Fabaceae-infecting orthotospoviruses (de Oliveira et al., 2012 and Zhou and Tzanetakis 2019). Episodic diversifying selection evidence was found between BeNMV and SVNV only in the NSm and RdRp (de Oliveira et al., 2012). Interestingly, BeNMV-NSm was shown to be inefficient in systemically transporting Alfalfa mosaic virus (AMV) in infiltrated plants compared to other orthotospoviruses (TSWV and Tomato chlorotic spot virus (TCSV)) (Leastro et al., 2017). As we were unable to localize the NSm from SVNV for comparison, it remains to be seen if localization to the plasmodesmata was impaired, but suggests severe challenges to movement in SVNV consistent with what was seen in BeNMV.

This lethality of SVNV-NSm may also be responsible for the inability of SVNV to move systematically on its own in soybeans. The SVNV-NSm belongs to the ‘30K’ movement protein superfamily, distinguished by the presence of the highly conserved LxDx40G motif (Zhou and Tzanetakis 2019) implicated in cell-to-cell movement (Mushegian and Koonin 1993 and Silva et al., 2001). However, a notable substitution was observed at the beginning of this motif, where the Leu residue was replaced by an Ile (Zhou and Tzanetakis 2019). Unlike some orthotospoviruses such as Tomato spotted wilt virus (TSWV) and Groundnut bud necrosis virus (GBNV), SVNV-NSm lacks the “P/D-L-X motif” and phospholipase A2 catalytic sites (Silva et al., 2001 and Kormelink 1992). This discrepancy may explain why SVNV-NSm induces cell death as important motifs are absent for the tubule formation required for the functionality of SVNV-NSm (Zhou and Tzanetakis 2019). NSm of TSWV triggered TSWV-like symptoms in *N. benthamiana*, and its constitutive expression in transgenic *N. tabacum* is sufficient to cause severe, infection-like symptoms (Prins et al., 1997, Lewandowski and Adkins, 2005, Li et al., 2009, and Rinne et al., 2005). The SVNV-NSm lethality could explain symptoms of necrosis in soybeans. However, NSm was able to be expressed in insect cells, which may indicate that these substitutions or absent amino acids are needed in plant cells to express but may be not crucial in insect cells. We hypothesize that thrips are a better host for SVNV and the ability to spread and move throughout the US may be due to the insect rather than success in plants.

The mutations that we detected in our SVNV-NSm sequences from the field were not found to be at any of the reported motifs suggesting that these motifs are highly conserved among orthotospoviruses (Li et al., 2009). The three detected mutations in NSm were the least conserved among the three sequenced proteins (Figure 6-C). This indicates that SVNV-NSm has the lowest diversity among the sequenced three proteins and changes the least. SVNV-NSm infiltrated alone did not express and was toxic to infiltrated leaves, however, in the sequencing study, we were able to extract, clone, and sequence this gene from naturally infected soybean plants in Alabama. This may indicate that the presence of all SVNV proteins during infection may stabilize the cell conditions for NSm expression or that the RNA is produced successfully, but the protein is toxic (Chen and Zacharias 2023). Moreover, since SVNV systemic movement during infection in soybean was linked to the presence of Bean pod mottle virus (BPMV) (Zhou and Tzanetakis 2020), maybe the presence of a helper virus could allow the virus to move in conjunction with the movement protein with BPMV. Additionally, the mechanical inoculation *N. benthamiana* leaves from SVNV infected plants showed that SVNV was detectable in the top and lower non-inoculated leaves (Data not shown), which indicated that SVNV was able to systemically move to and replicate in the new leaves in other hosts. These results may indicate that SVNV-NSm strains from AL may express if they were localized or that spread in *N. benthamiana* is not linked to NSm specifically.

The glycoprotein, G_N_, localized in the cell and nuclear membrane in INSV, IYSV, and TSWV but for CaCV, where it localized in the cell nucleus (Dietzgen et al., 2012, Tripathi et al., 2015, Widana Gamage and Dietzgen 2017, and Martin et al., 2024). SVNV-G_N_ in both plants and insects was localized to the cell membrane and around the cell nucleus. Additionally, in insects, the localization of G_N_ appeared much more cytoplasmic than ever seen with TSWV-G_N_ (Martin et al., 2024). Moreover, the soluble form of G_N_ (G_N_-S) formed aggregates in insect cells which was not expected as Martin et al., noted that these aggregates were not seen forming with TSWV-G_N_-S (Martin et al., 2024). The glycoprotein, G_C,_ in plants is localized to cell membranes in TSWV and to the cell and nuclear membrane in INSV, IYSV, and CaCV. SVNV-G_C_ in plants was also localized to the cell and nuclear membrane similar to TSWV-G_C_ (Martin et al., 2024). There was no detectable localization of G_C_ in insect cells which is similar to that seen previously with TSWV and may be due to potential toxicity in insect cells or the large size of the protein (Martin et al., 2024). Based on these protein localization results, SVNV has shown a distinct localization pattern compared to other discussed orthotospoviruses, especially, for its N and NSs that localized to the cell nucleus contradicting all characterized orthotospoviruses except CaCV-NSs (Widana Gamage and Dietzgen 2017), and NSm being lethal to plant cells. This is consistent with our hypothesis based on the phylogenetic tree.

The unique association between SVNV-N and NSs over the six days’ time course (Supplementary Figure 2-C: XX) suggests that these two proteins may interact as shown in CaCV (Widana Gamage and Dietzgen 2017). Furthermore, the accumulation of both proteins outside and around the nucleus (Supplementary Figure 2-B: IX-XX) which formed voids may indicate that the replication of SVNV might take place in regions around the nucleus. This is supported by the localization of NSs and N when expressed alone inside the nucleus. This suggests that SVNV may have a closer association to the nucleus or nuclear proteins than other previously characterized orthotospoviruses. These observations warrant future studies to directly look at protein-protein interactions and protein localizations during infection.

When SVNV was first reported in AL, it was confirmed by sequencing 250 nucleotides in the RNA dependent RNA polymerase (RdRp) (Accession number: KC431808), and no sequencing was conducted on any other genes. Although, the RdRp is conserved and this enabled an identification of the virus, it did not allow a detailed comparison to the reference strain. Comparing these mutations to that from a thrips sample shows that the mutations are present in both plants and the insect suggesting these mutations are in mature virions acquired by the insect and not simply part of a viral cloud. In the protein phylogeny of N and NSs, the SVNV TN sequence was found closely related to the northern SVNV samples possibly due AL bordering TN in the north. In the two phylogenic trees of NSs and NSm, the central AL sequences blended with northern sequences (Figure 7-B and C). Additionally, the sequences of N and NSm that were amplified from thrips were found to be grouped with northern sequences, however, the thrips were collected from central AL. This may indicate that SVNV moves from northern to central AL with the thrips carrying the virus. Finally, the previous diversity studies conducted by (Zhou and Tzanetakis 2013) indicated that SVNV has a relatively homogeneous population structure within the geographic states studied. When we added our sequences to their sequences, we found that the diversity of SVNV increased by 16% for the amino acid sequences.

## Conclusions

Here we introduce the localization behavior of this important orthotospovirus in both plant and insect cells supporting our hypothesis that they are unique to this system. This revealed key findings as the N and NSs localized to both the cytoplasm and the nucleus, which is a unique finding for N compared to other systems. Additionally, NSm was lethal to cells when infiltrated into plants, but not when transfected into insect cells. We also demonstrated the second part of our hypothesis that the sequence of SVNV has undergone changes since the original sequencing study was conducted over ten years ago. SVNV ORFs for N, NSs and NSm have never been fully sequenced from AL, and in this study, we provided 88 SVNV sequences that revealed conserved and non-conserved aa mutations. Mutations from SVNV for these ORFs is conserved from insects when compared to the plant samples, suggesting that the mutations are acquired by insects as part of a mature virion. Furthermore, it is clear that the diversity of SVNV is increasing which could correlate to the increase in viral incidence in Alabama (Sikora et al., 2018) and it is necessary to further explore these changes in the full genome. This will help us gain knowledge about how SVNV evolves which will help us to understand what we might expect from SVNV in the future.

---

**Supplementary Table 1:** Percentage of the natural amino acid mutations observed in N, NSs, and NSm genes of Soybean vein necrosis virus collected from infected soybean fields from Alabama and from infected thrips collected from AL.

| No | SVNV-N protein (32 samples*) |  |  | SVNV-NSs protein (25 samples*) |  | SVNV-NSm protein (31 samples *) |  |  |
| --- | --- | --- | --- | --- | --- | --- | --- | --- |
|  | Amino acid mutations and their site | Mutation percentage | Mutation detected in Thrips? | Amino acid mutations and their site | Mutation percentage | Amino acid mutations and their site | Mutation percentage | Mutation detected in Thrips? |
| 1 | A13V | 93.75 % | Yes | S34N | 21 = 84 % | S19G | 25.80 % | No |
| 2 | I15L | 100 % | Yes | I64V | 21 = 84 % | S63N | 48.38 % | Yes |
| 3 | K22R | 90.62 % | Yes | M261V | 25 = 100 % | V143I | 41.93 % | Yes |
| 4 | A115E | 87.5 % | Yes | R303K | 25 = 100 % |  |  |  |
| 5 | D118E | 59.37 % | Yes | V310I | 25 = 100 % |  |  |  |
| 6 | L128F | 56.25 % | Yes | L334I | 21 = 84 % |  |  |  |
| 7 | G169N | 84.37 % | Yes |  |  |  |  |  |
| 8 | I227M | 68.75 % | Yes |  |  |  |  |  |

