## Supplementary figures and images for "Characterization of Soybean Vein Necrosis Virus (SVNV) Proteins: Sequence Analysis of Field Strains and Comparison of Localization Patterns in Differing Cell Types"

### Supplemental Figure 2

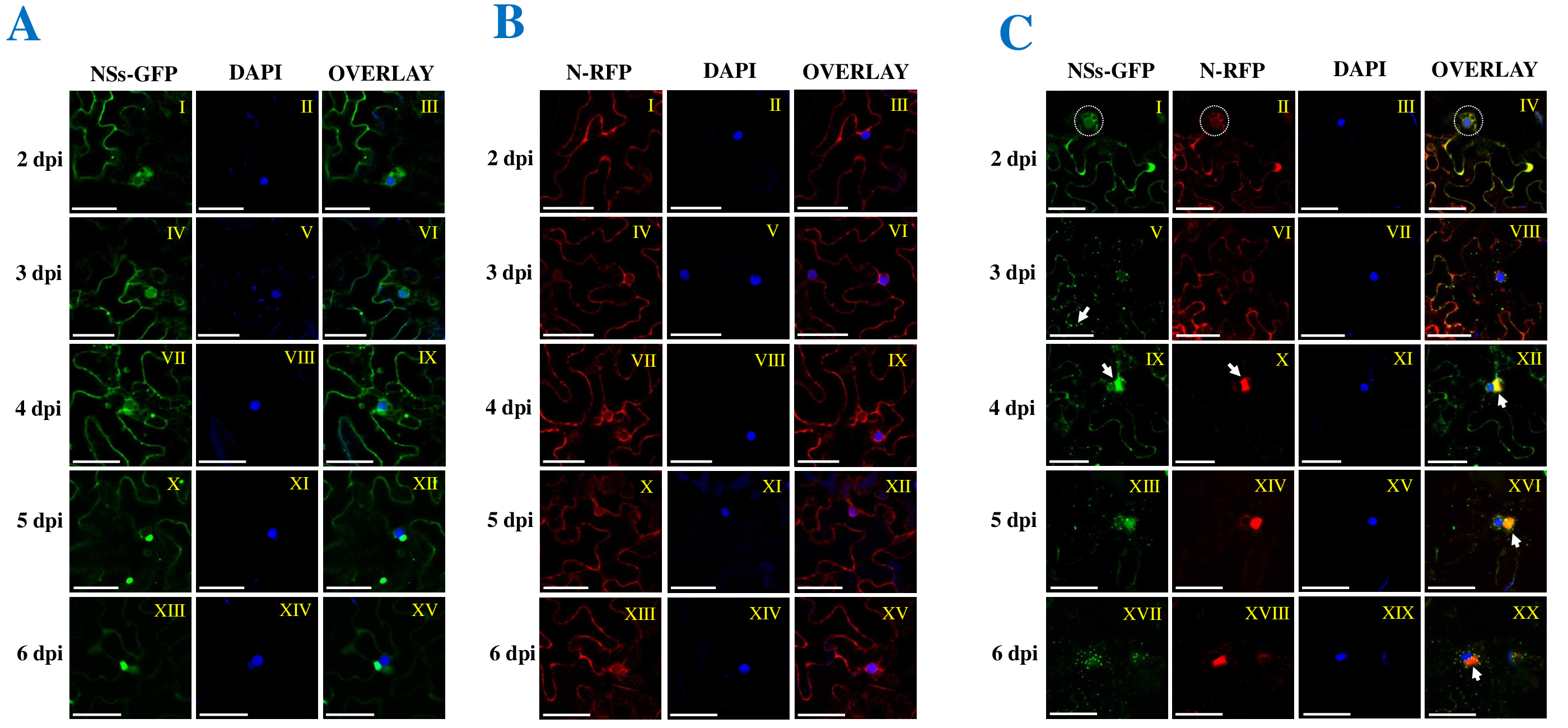

### Supplementary Figure 1

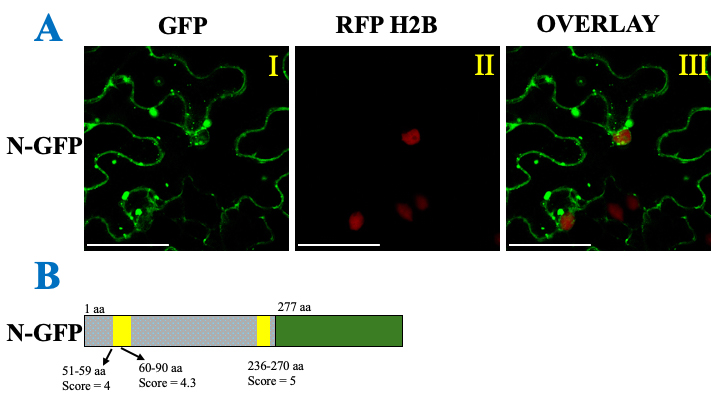
